# *Wolbachia*-induced Cytoplasmic Incompatibility drives epigenetic and maternally-influenced post-embryonic defects

**DOI:** 10.64898/2026.04.15.718768

**Authors:** Claire Perez, Jillian Porter, Brandt Warecki, William Sullivan

## Abstract

A common form of *Wolbachia*-induced manipulation of host reproduction is Cytoplasmic Incompatibility (CI). In CI, *Wolbachia* modification of sperm results in pronounced defects in paternal chromosome condensation, replication, and segregation during the first mitotic division. Recent studies in *D. simulans* demonstrate that CI also induces independent and distinct later developmental defects resulting in high rates of mitotic errors during the mid-blastula transition and larval lethality. Here we show that in *D. melanogaster*, embryos derived from CI crosses experienced significant mitotic defects during gastrulation and increased larval lethality, both of which were eliminated in the progeny of Rescue crosses (both sexes infected). Examination of CI using females from 13 genetically distinct wild-type lines of the *Drosophila* Genetic Reference Panel (DGRP) revealed significant variation in the strength of the CI-induced lethality. Early embryonic pre-hatching and late larval lethal phases were uncorrelated, suggesting distinct factors influence the extent of the two lethal phases. Additionally, 3^rd^ instar larvae and adults derived from *D. melanogaster* CI crosses exhibited locomotor defects that were also eliminated in Rescue crosses. These studies support a model in which *Wolbachia* effects on the sperm chromatin produce delayed developmental and locomotor defects, suggesting the involvement of epigenetic mechanisms. Support for this idea comes from our finding that levels of the heritable chromatin mark H3K27me1 are significantly elevated in CI-derived embryos. We conclude that the full measure of CI strength should take into account pre- and post-hatching lethality as well as locomotor defects. Together our findings suggest that the strength of these CI-induced phenotypes is governed at least in part by epigenetics and the maternal genetic background.

**AUTHOR SUMMARY:** Since the discovery of the antiviral properties of the bacteria *Wolbachia*, numerous strategies using this insect endosymbiont have been developed to combat vector-borne disease. While the success of these strategies relies on the rapid spread of *Wolbachia* through a naturally uninfected insect population, the molecular mechanisms by which *Wolbachia* promote their spread remain poorly defined. Current research on the primary mechanism behind *Wolbachia* spread, cytoplasmic incompatibility (CI), focuses on understanding the dramatic decrease in egg hatch rates that occurs when uninfected females mate with infected males. Here, we demonstrate that CI also induces substantial post-hatching larva and adult locomotor defects and lethality. In accord with these developmentally delayed defects, we show *Wolbachia* dramatically alter an epigenetic chromatin mark. Finally, we show that host maternal factors contribute to CI strength. Taken together, these results demonstrate that CI induces a much more expansive and developmentally delayed suite of phenotypes than previously reported.

## INTRODUCTION

*Wolbachia* are a widespread bacterial endosymbiont present in the germline of many evolutionary diverse insect species (1, 2). Like mitochondria, these bacteria are transmitted through the female germline to all of their progeny. Consequently, in many insect species, *Wolbachia* modify host reproduction such that infected females are favored over uninfected females. Modifications include transformation of males into fertile females, male killing, induction of parthenogenesis, and cytoplasmic incompatibility (CI) (3). The latter is unique among these reproductive modifications as it involves a collaborative action of *Wolbachia* in both sexes. Crosses between uninfected females and infected males result in a dramatic reduction in egg hatch rates (the CI cross) (4, 5). In contrast, crosses between infected females and either infected males (the Rescue cross) or uninfected males (the reciprocal cross) yield normal egg hatch rates. In populations with high *Wolbachia* infection frequencies, infected females enjoy a selective advantage over uninfected females, contributing to their rapid spread through host populations (6, 7). Application of *Wolbachia*-mediated CI has proven an efficient strategy in the management of pest and disease-bearing insects (8, 9).

Since the discovery of this phenomenon, a major goal has been to elucidate the molecular and cellular basis of the *Wolbachia*-induced CI phenotypes, as well as its suppression in Rescue crosses (10). Studies in *D. simulans* and other species demonstrate that segregation failures of the paternal chromosomes during the first mitotic division are a primary cause of failed egg hatch (11, 12, 13, 14, 15). Subsequent studies revealed earlier defects in the protamine-to-histone transition, incomplete DNA replication, and failed chromosome condensation (16, 17, 18, 19). Insight into *Wolbachia* effector proteins responsible for inducing CI and Rescue has come from identifying a pair of *Wolbachia* prophage-embedded genes, known as *cifs* (Cytoplasmic Incompatibility factors). The *cif* gene, *cidB/cinB,* when ectopically expressed either singly or together with its cognate *cidA/cinA* in the *D. melanogaster* male germline, produces equivalent first division chromosome defects and reduced egg hatch (19, 20, 21, 22). Strikingly, when crossed to females ectopically expressing the “A” component alone, the first zygotic division occurs normally (19, 20, 22, 23, 24, 25, 26).

While CI-induced lethality has been extensively studied by focusing on first division paternal chromosome segregation defects and embryonic lethality, less attention has been devoted to CI-induced lethality that occurs after embryo hatching during larval and pupal stages. Studies in the haplodiploid species *P. kellyanus* revealed CI crosses generate significant post-embryonic lethality (27). Similarly, in *D. simulans*, a portion of progeny derived from CI crosses die during the larval stages following hatching (28) (Fig 1). Significantly, this late lethality is eliminated in Rescue crosses. In addition, a large fraction of *D. simulans* CI-derived embryos progress normally through the initial syncytial divisions only to experience extensive segregation errors during cellularization and the mid-blastula transition (28). Furthermore, in *D. melanogaster,* wMel and *cif*-transgene expression yields post-first division defects that may be rescued by *cifA* in the female (20, 29). Whether these segregation errors and late embryonic defects contribute to post-hatching lethality is unclear. Regardless, these studies demonstrate that the *Wolbachia*-induced defects on the paternal chromosome complement produce deleterious, developmentally delayed effects through an unknown mechanism. Studies demonstrating *Wolbachia* induce altered patterns of histone methylation on the paternally derived chromosomes suggest consequential epigenetic chromatin marks may be responsible for at least some of the CI-induced phenotypes (30).

**Fig 1.**
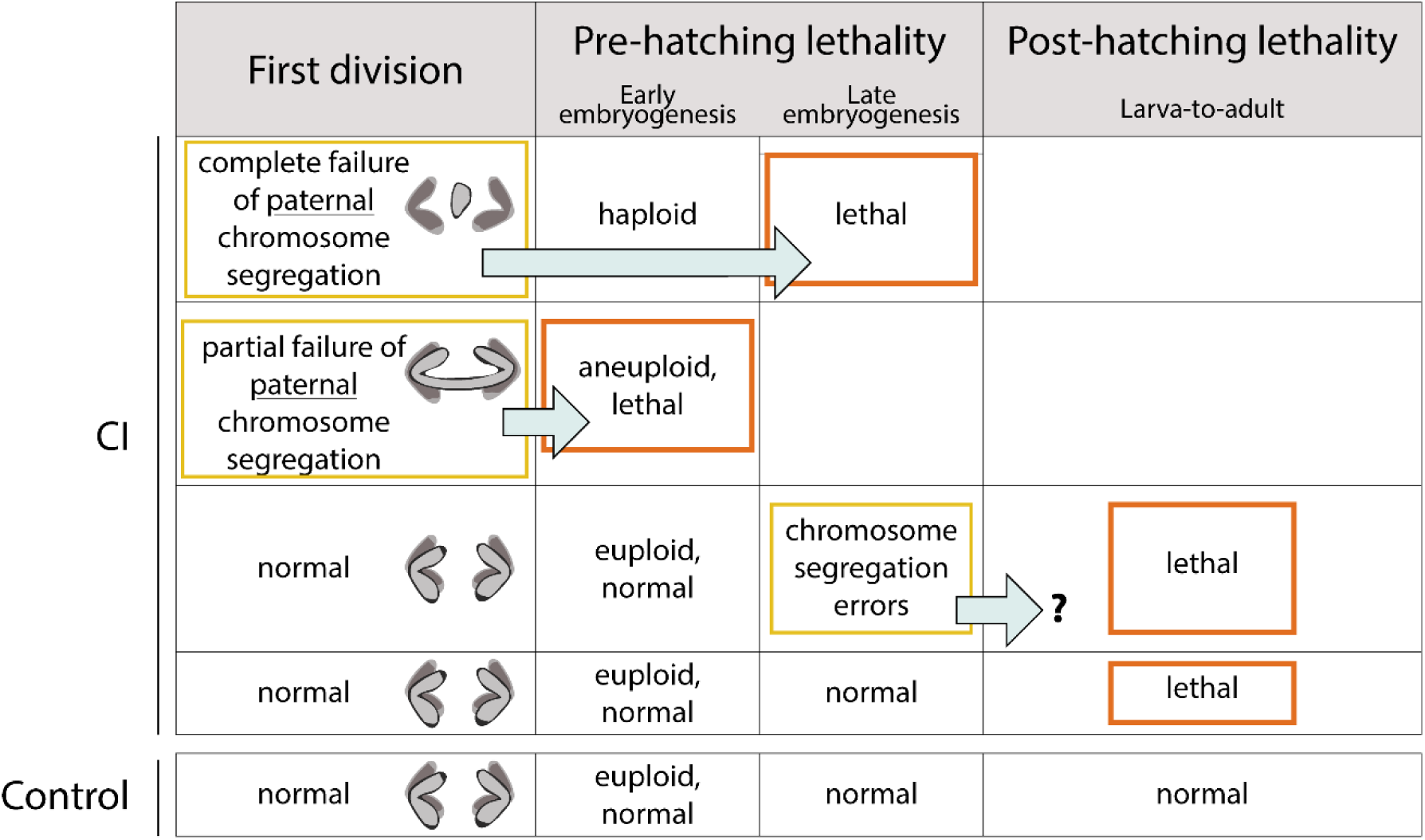
CI-derived progeny experience pre- and post-hatching lethality. During the first division, CI-derived embryos exhibit either normal, partial, or complete paternal chromosome segregation errors. Partial paternal chromosome segregation errors produce aneuploidy and early embryonic lethality. Complete segregation errors produce haploid embryos that die in late embryogenesis prior to hatching. A fraction CI-derived embryos that progress normally through the first and subsequent divisions undergo segregation errors during gastrulation. This may contribute to the observed post-hatching larva-to-adult lethality. Finally, there is a class of CI-derived embryos that develop normally through embryogenesis and hatch but die during larval to adult development.

Here we investigated whether the CI-induced developmentally delayed defects observed in *D. simulans* are a conserved property of CI by examining this issue in *D. melanogaster*. We found that in *D. melanogaster*, like *D. simulans,* CI also produced extensive embryonic chromosome segregation defects as well as post-hatching larval developmental defects. As with *D. simulans*, we found many of these CI-induced defects were developmentally delayed. We also discovered that 3^rd^ instar larvae and adults derived from *Drosophila* CI crosses suffer from behavioral/locomotor defects. To ensure our studies are relevant to populations with genetically diverse backgrounds, we chose to work with the well-characterized *Drosophila* Genetic Reference Panel (DGRP). This is a set of highly inbred, sequenced lines isolated from Raleigh, North Carolina in 2003 (31). By crossing uninfected DGRP females to OreR wMel-infected males, we demonstrated that the maternal genetic background dramatically influences the strength of CI-induced lethality.

Collectively, these early, late, and behavioral phenotypes were eliminated in Rescue crosses. Our finding that H3K27me1 levels were significantly elevated in gastrulating CI-derived embryos supports the involvement of epigenetic mechanisms in delayed CI-induced phenotypes.

## RESULTS

### CI-derived *D. melanogaster* embryos exhibit increased mitotic errors during gastrulation

Although classic CI-induced first embryonic division defects are well documented, some embryos escape these defects and develop to later embryonic stages. Previous studies in *D. simulans* demonstrated that these CI “escapers” progress normally through the interior syncytial nuclear divisions (nuclear cycles 1-9) before experiencing significant division errors during the late cortical divisions (nuclear cycles 12-14) and gastrulation (28). Because CI is generally weaker in *D. melanogaster* than in *D. simulans*, we wished to determine if late-developing CI-derived embryos also experienced delayed chromosome segregation errors in *D. melanogaster*. First, to gauge CI strength in our crosses for these experiments, we concomitantly conducted egg hatch (egg-to-larva) assays for Control, CI, and Rescue crosses (Fig 2A). 96% +/− 3% of Control eggs, 76% +/− 10% of CI eggs, and 99% +/− 3% of Rescue eggs hatched. Adjusted pairwise comparisons demonstrated that CI egg hatch was significantly lower than either Control (*p* = 0.007) or Rescue (*p* = 0.004) egg hatches, with no significant difference in egg hatch between Control and Rescue crosses (*p* = 1). Thus, we observed a mild, but significant CI in our experimental conditions, with respect to egg hatch rates.

**Fig 2.**
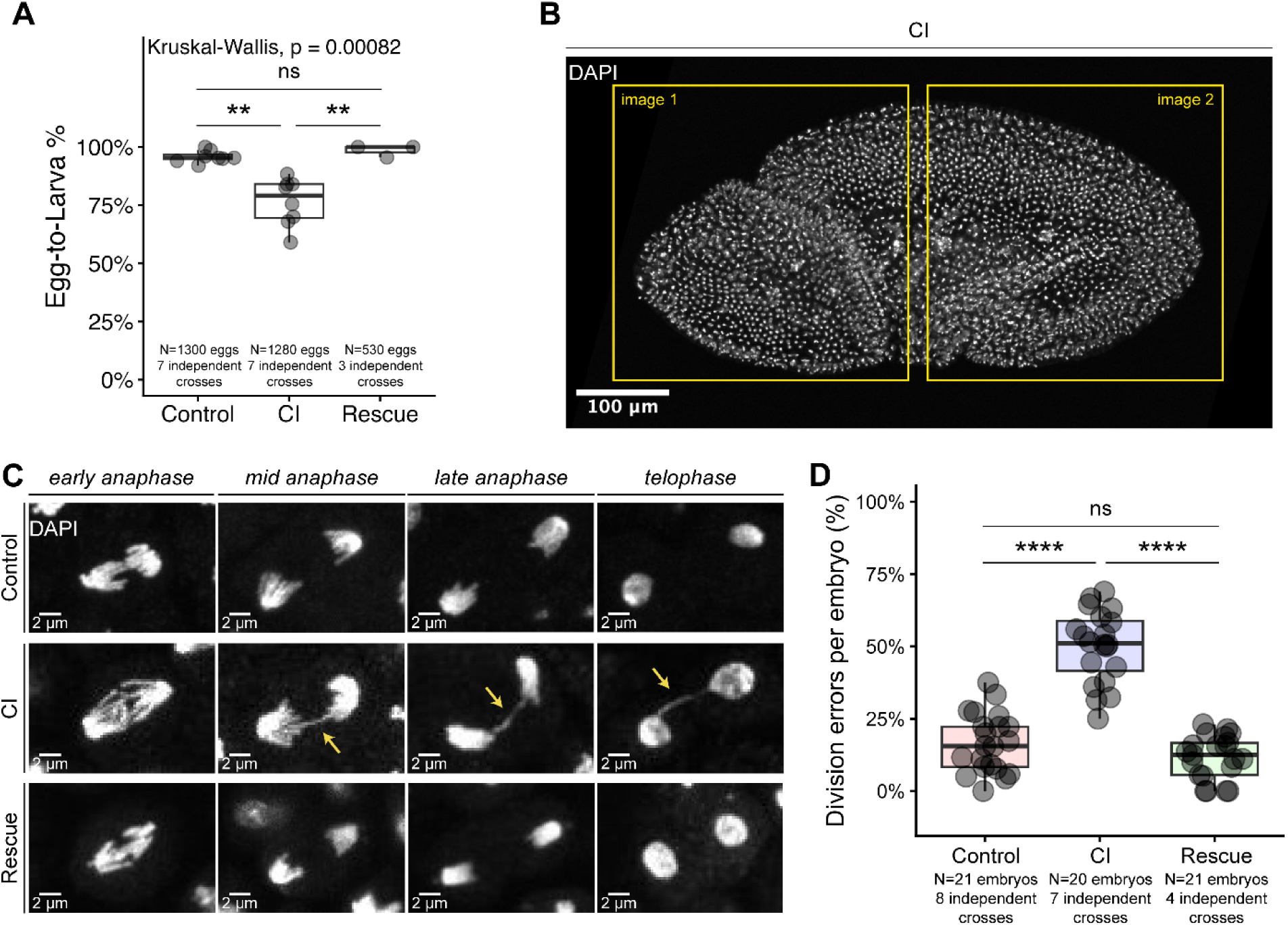
CI-derived *D. melanogaster* embryos undergo mitotic defects during gastrulation. **A)** Graph quantifying measured egg hatch for Control, CI, and Rescue crosses used to assay late embryonic chromosome segregation errors. **B)** A DAPI-stained gastrulating embryo. For assaying chromosome segregation errors, each embryo was imaged twice at high magnification (to cover a substantial volume of the embryo) and scored for the percentage of mitotic cells that exhibited chromosome segregation errors. The boxes indicate approximate areas where embryos were imaged. **C)** Images of dividing nuclei in early, mid, and late anaphase and telophase in Control-derived (top row), CI-derived (middle row), and Rescue-derived (bottom row) gastrulating embryos. **D)** Graph quantifying the mitotic error rates per embryo during gastrulation. Statistical significance was assessed by a Dunn’s test using the Bonferroni correction to adjust for multiple comparisons **(A)** or using a linear mixed model, with pairwise comparisons of estimated marginal means performed in the emmeans package in R, adjusting for multiple comparisons with the Tukey method **(D)** **** = *p* < 0.0001; ** = *p* < 0.01; ns = *p* > 0.05.

Immediately following cellularization during interphase of nuclear cycle 14, embryos undergo gastrulation. During gastrulation, rounds of cell division occur bilaterally in specific regions of the embryo known as mitotic patches (32, 33). In CI-derived *D. simulans* embryos, mitotic patches experience significantly increased segregation errors (28). To determine if CI-derived *D. melanogaster* embryos also exhibit increased segregation errors at this stage, we scored mitotic cells in gastrulating embryos derived from Control, CI, and Rescue crosses (Fig 2B). We observed multiple examples of division errors in CI-derived gastrulae, including anaphases with bridged and lagging chromosomes (Fig 2C). Control-derived gastrulating embryos underwent chromosome segregation errors in 17% +/− 10% of divisions, compared to 50% +/− 13% of divisions from CI-derived embryos and 12% +/− 8% of divisions in Rescue-derived embryos (Fig 2D). Adjusted pairwise comparisons revealed that CI-derived gastrulating embryos experienced significantly higher rates of errors than both Control-derived embryos (CI-Control 95%CI [0.26,0.42], *p* = 1.8×10^−11^) and Rescue-derived embryos (CI-Rescue 95%CI [0.24,0.51], *p* = 1.71×10^−4^), while there was no significant difference in the amount of division errors between Control-derived and Rescue-derived embryos (Control-Rescue 95%CI [−0.10,0.17], *p* = 0.74). These results highlight that CI causes developmentally delayed defects in chromosome segregation during the gastrulae stage of *D. melanogaster* embryos, similar to those which are observed in *D. simulans*.

### The chromatin of CI-derived embryos maintains increased levels of methylation

Our results demonstrate that embryos derived from CI crosses that escaped first division defects suffer chromosome segregation errors during gastrulation. These results suggest that the CI-induced disruptions to the paternal chromosomes may be epigenetic in nature. Support for this idea comes the findings in *Nasonia vitripennis* that show dramatic alterations in the histone methylation levels on the paternal chromatin of first division CI-derived embryos (30). However, whether a similar increase in histone methylation occurs in *Drosophila* has not been determined. Furthermore, it is unknown whether the CI-induced increased methylation can be maintained and transmitted through multiple rounds of mitosis.

To address these questions, we used immunofluorescent analysis to examine H3K27me1 levels in *D. melanogaster* Control- (Fig 3A), CI- (Fig 3B), and Rescue- (Fig 3C) derived gastrulating embryos. Surface level views of the gastrula appeared to show brighter H3K27me1 signal in CI-derived embryos compared to Control- or Rescue-derived embryos, although several nuclei in Rescue-derived embryos were sometimes bright with H3K27me1 staining and overall staining of Rescue-derived embryos was variable (Fig 3A-C). To quantify any difference in H3K27me1 levels between Control-, CI-, and Rescue-derived embryos, we imaged gastrulae at the midplane and calculated the H3K27me1/DAPI signal intensity ratio corresponding to cells at the edge of the embryos (Fig 3D). To control for effects between different experimental replicates, we normalized all values from that replicate to the average value of the Control-derived embryos for that replicate. Control-derived gastrulating embryos had a normalized ratio of H3K27me1/DAPI of 1 +/− 0.23. CI-derived gastrulating embryos had a normalized ratio of H3K27me1/DAPI of 1.35 +/− 0.37. Rescue-derived gastrulating embryos had a normalized ratio of H3K27me1/DAPI of 0.87 +/− 0.32 (Fig 3E). Adjusted pairwise comparisons revealed that CI-derived gastrulating embryos experienced significantly higher ratios of H3K27me1/DAPI than both Control-derived embryos (CI/Control 95%CI [1.16,1.40], *p* = 1.2×10^−8^) and Rescue-derived embryos (CI/Rescue 95%CI [1.30,1.57], *p* = 2.1×10^−14^), while Control-derived embryos had a higher H3K27me1/DAPI ratio than Rescue-derived embryos (Control/Rescue 95%CI [1.02,1.23], *p* = 0.02). Thus, like in *Nasonia*, H3K27me1 levels are increased in CI-derived embryos in *D. melanogaster*. Importantly, this increase is maintained through multiple rounds of cell division.

**Fig 3.**
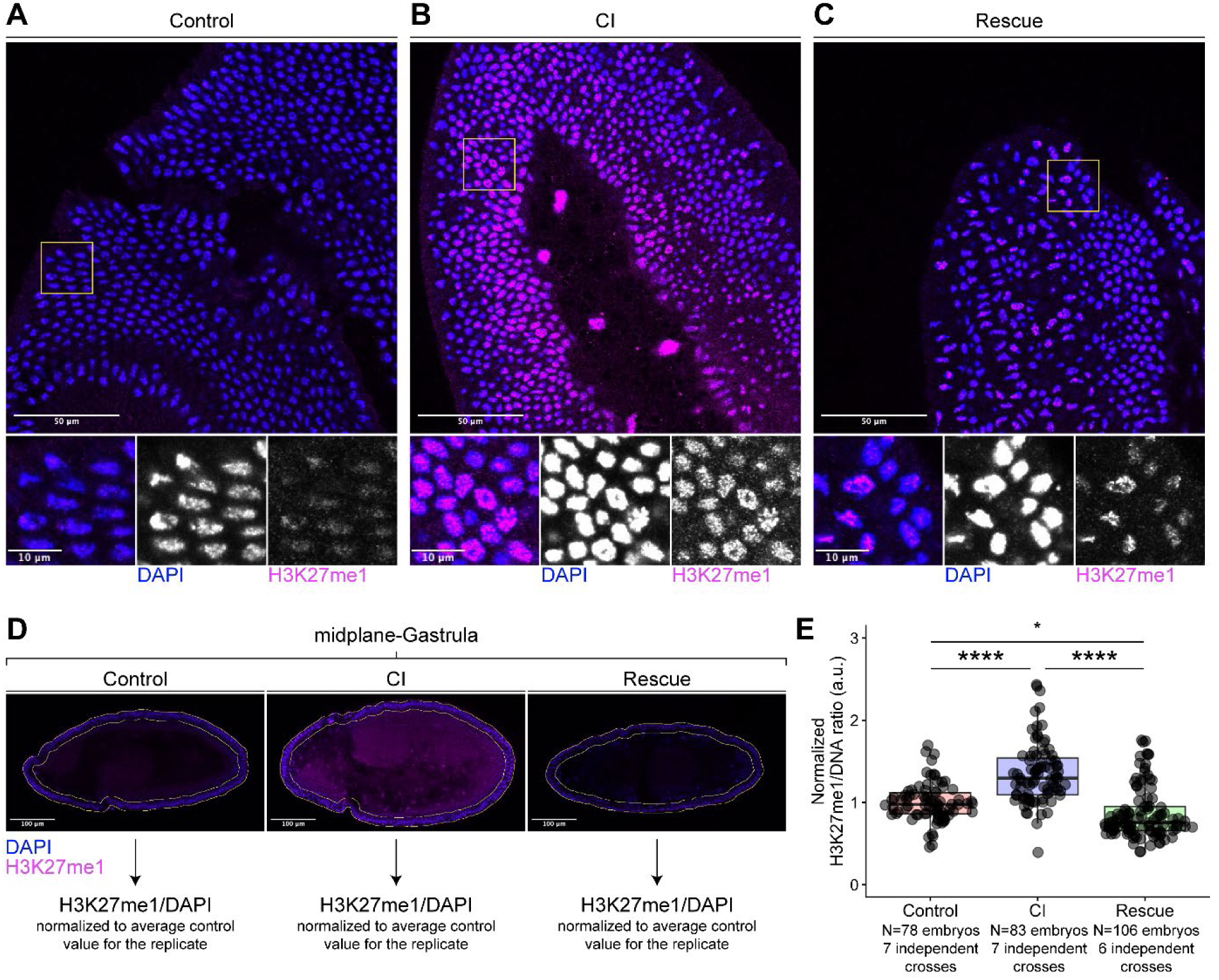
H3K27me1 levels are enhanced in *D. melanogaster* CI-derived embryos. **(A-C)** Control-derived **(A)**, CI-derived **(B)**, and Rescue-derived **(C)** gastrulating embryos were fixed and double stained with two chromatin markers: DAPI and anti-H3K27me1. **D)** Embryos were scored by taking an image of the entire embryo at the midplane. DAPI and H3K27me1 intensities were measured in a region of interest consisting of a 20 µm band around the edge of the embryo (where cells are in cross section). The H3K7me1/DAPI ratio was calculated for each embryo and normalized to the average H3K27me1/DAPI ratio of the Control embryos scored for that independent replicate. **E)** Graphs quantifying the measured ratios of normalized H3K27me1/DAPI per embryo. Statistical significance was assessed using a generalized linear mixed model, with pairwise comparisons of estimated marginal means performed in the emmeans package in R, adjusting for multiple comparisons with the Tukey method **(E)** **** = *p* < 0.0001; * = *p* < 0.05..

### Larvae and adults derived from *D. melanogaster* CI crosses exhibit climbing defects

Given these results, we wished to determine if CI-derived progeny experienced any post-hatching defects as well. Although 3^rd^ instar larvae derived from CI crosses appeared normal, we observed that they did not climb as high as larvae derived from Control or Rescue crosses in preparation for pupation. To quantify this phenotype, we scored the vertical positions of pupae as either above or below a physical rim at the neck of the bottle, 2 cm above the media (Fig 4A-B). On average, 53% +/− 17% of Control-derived larvae pupated above the 2 cm threshold compared to 20% +/− 26% of CI-derived larvae and 57% +/− 20% of Rescue-derived larvae (Fig 4C). Adjusted pairwise comparisons demonstrated that CI-derived larvae pupated at a significantly lower rate above the 2 cm threshold than either Control-derived (CI/Control 95%CI [0.15,0.74], *p* = 0.004) or Rescue-derived larvae (CI/Rescue 95%CI [0.11,0.62], *p* = 0.0008), while there was no significant difference in the percentages of larvae pupated above 2 cm between Control and Rescue offspring (Control/Rescue 95%CI [0.45,1.45], *p* = 0.67). To compare the strength of this phenotype to the strength of CI-induced embryonic lethality, we also performed egg hatch assays from these crosses (Fig 4D). 96% +/− 4% of Control eggs, 79% +/− 8% of CI eggs, and 97% +/− 2% of Rescue eggs hatched. Adjusted pairwise comparisons demonstrated that CI egg hatch was significantly lower than either Control (CI/Control 95%CI [0.12,0.27], *p* = 3.5×10^−14^) or Rescue egg hatches (CI/Rescue 95%CI [0.10,0.24], *p* = 2.8×10^−14^), with no significant difference in egg hatch between Control and Rescue crosses (Control/Rescue 95%CI [0.51,1.41], *p* = 0.73). Thus, even with a relatively “weak” CI according to egg hatch, CI-derived offspring exhibited a dramatic CI-induced phenotype during their larval and pupal stages.

**Fig 4.**
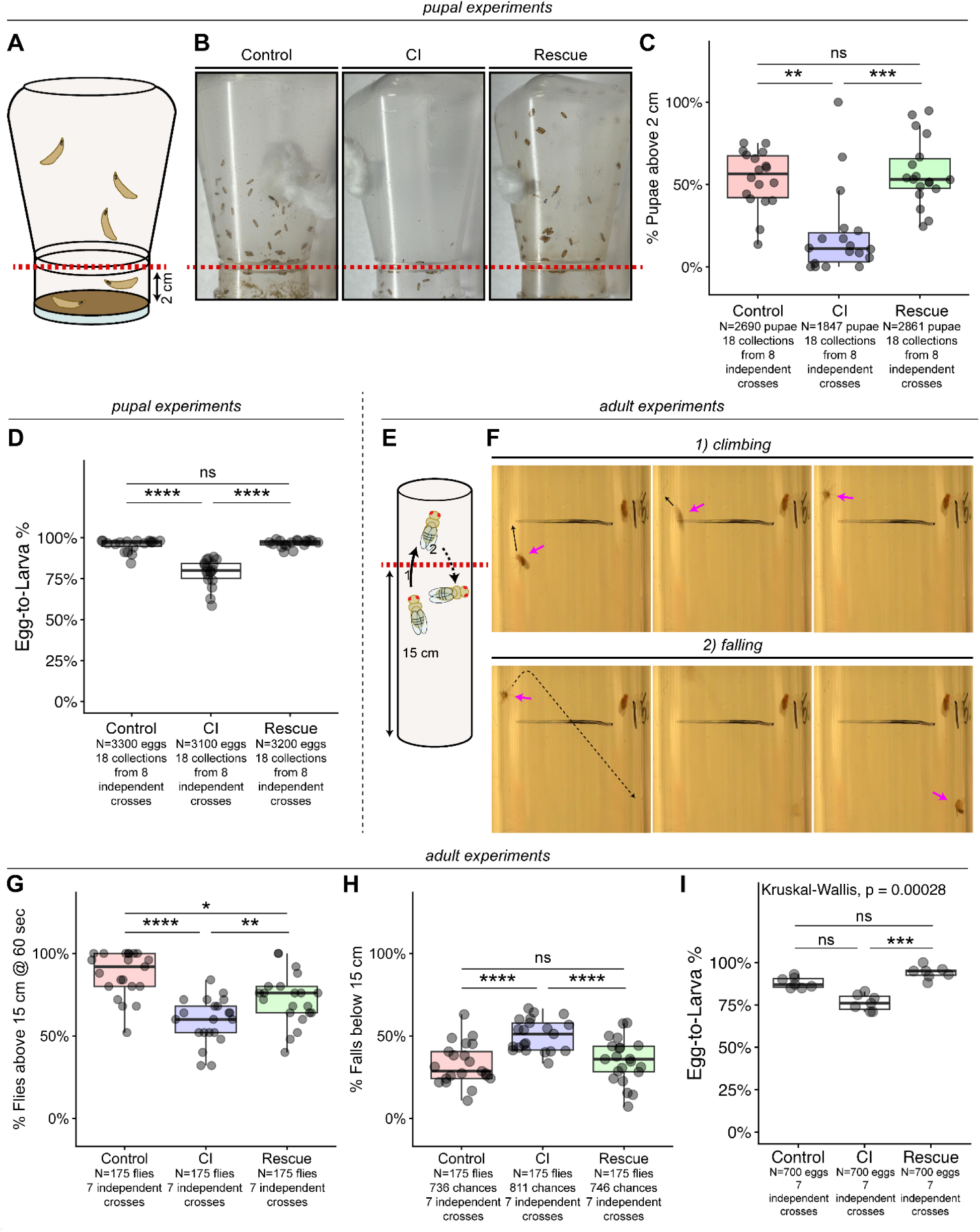
Larvae and adults derived from *D. melanogaster* CI crosses exhibit locomotor defects. **A)** The percentage of larvae that pupated 2 cm above the media was measured for each cross. At 2 cm above the media, there was also a phsyical rim in the neck of the bottle that larvae had to traverse. **B)** Representative images of the pupation heights of Control-derived, CI-derived, and Rescue-derived progeny. Dashed red line indicates the 2 cm height threshold. **C)** Comparison of the percentage of pupae above the 2 cm line in each cross. **D)** Graph quantifying measured egg hatch for Control, CI, and Rescue crosses used to assay pupation heights. **E)** We measured the percentage of adult flies that climbed above a 15 cm threshold. The recordings were focussed on the 15 cm line. To avoid double counting, we counted 1) the number of flies that had climbed above 15 cm and then subtracted 2) the number of those flies that then fell or crawled below 15 cm. **F)** An example of one CI-derived fly (pink arrows) climbing above 15 cm and then falling below. **G)** Comparision of the percentage of flies that climbed and remained above 15 cm at the end of the assay. **H)** Comparison of the percentage of flies that had been above 15 cm but then fell below 15 cm threshold throughout the entire time of the assay. **I)** Graph quantifying measured egg hatch for Control, CI, and Rescue crosses used to assay adult locomotor defects. Statistical significance was assessed by a Dunn’s test using the Bonferroni correction to adjust for multiple comparisons **(I)** or using linear (**H**) or generalized (**C,D,G**) linear mixed models, with pairwise comparisons of estimated marginal means performed in the emmeans package in R, adjusting for multiple comparisons with the Tukey method **** = *p* < 0.0001; *** = *p* < 0.001; ** = *p* < 0.01; * = *p* < 0.05; ns = *p* > 0.05.

As the CI-derived larvae have normal morphology, we suspect these locomotor defects are neurologically based. These results clearly demonstrate that *Wolbachia* induces developmentally delayed post-hatching defects in the progeny of CI crosses. This finding was in accord with a model in which some of the CI-induced phenotypes are the result of epigenetic chromatin modifications. Additionally, like the *Wolbachia*-induced embryonic mitotic division defects, these larval locomotor defects are ameliorated in the Rescue cross.

These results motivated us to test whether adults derived from CI crosses also exhibited locomotor defects. We performed a negative-geotaxis assay with adults derived from Control, CI, and Rescue crosses and scored for the number of flies above 15 cm within 60 seconds (Fig 4E-F). At the end of the assay, 87% +/− 14% of Control-derived adults were above 15 cm, compared with 59% +/− 14% of CI-derived adults and 72% +/− 15% of Rescue-derived adults (Fig 4G). Adjusted pairwise comparisons revealed that CI-derived adults were above 15 cm at a significantly lower rate than Control-derived (CI/Control 95%CI [0.11,0.41], *p* = 1.1×10^−7^) and Rescue-derived adults (CI/Rescue 95%CI [0.32,0.86], *p* = 0.006), while Control-derived adults were above 15 cm significantly more than Rescue-derived adults (Control/Rescue 95%CI [1.18,5.23], *p* = 0.01). Analysis revealed this was the result of CI-derived flies continually falling off the wall throughout the assay, indicating impaired coordination (Fig 4F). Of the flies that had climbed above 15 cm, 33% +/− 13% of Control-derived adults, 50% +/− 10% of CI-derived adults, and 35% +/− 14% of Rescue-derived adults fell below the 15 cm threshold (Fig 4G). Adjusted pairwise comparisons revealed that CI-derived flies fell at significantly higher rates than Control-derived flies (CI-Control 95%CI [0.10,0.25], *p* = 6.0×10^−6^) and Rescue-derived flies (CI-Rescue 95%CI [0.07,0.23], *p* = 5.7×10^−5^), while Control-derived and Rescue-derived flies did not fall at a significantly different rate (Control-Rescue 95%CI [−0.10,0.058], *p* = 0.8). Together, these results suggest that developmental defects associated with CI persist into adulthood as a locomotor impairment.

To compare these phenotypes with traditional measures of CI strength, we additionally conducted egg hatch assays for the crosses used in these experiments and found 88% +/− 3% of Control eggs, 76% +/− 5% of CI eggs, and 94% +/− 4% of Rescue eggs hatched (Fig 4I). Adjusted pairwise comparisons demonstrated that CI egg hatch was significantly lower than Rescue (*p* = 0.0002) but not Control (*p* = 0.06) egg hatches, with no significant difference in egg hatch between Control and Rescue crosses (*p* = 0.3). Thus, even when CI is weak with no statistically significant lethality during egg hatch, locomotor impairments in CI escapers are detected.

### *Wolbachia*-induced CI lethality in *D. melanogaster* is the result of both embryonic and larval lethality

As described above, in *D. simulans*, CI induces independent, distinct embryonic and post-hatching larval lethality, both of which are suppressed in the Rescue crosses (28). To determine whether CI also induces post-hatching lethality in *D. melanogaster*, we examined hatch and adult survival rates of progeny from the *D. melanogaster* CI cross. In addition, we hypothesized that CI-induced post-hatching lethality could be a factor in determining CI strength in genetically diverse wild-type lines. Therefore, we performed this experiment with females from 13 distinct lines of the well-characterized *Drosophila* Genetic Reference Panel (DGRP) (31). We crossed DGRP females to *w*Mel infected or uninfected male flies (OreR). Six of the chosen DGRP lines were naturally infected with *Wolbachia*. Antibiotics were used to cure these lines, enabling us to perform both CI (cured females) and Rescue crosses (infected females).

We compared CI strength with regards to both embryonic and larval lethality for all DGRP lines amalgamated together (Fig 5A). 93% +/− 5% of Control-derived eggs hatched, while 77% +/− 11% of CI-derived and 96% +/− 2% of Rescue-derived eggs hatched. Adjusted pairwise comparisons demonstrated that CI egg hatch was significantly lower than either Control (*p* = 7×10^−9^) or Rescue (*p* = 2×10^−6^) egg hatches, with no significant difference in egg hatch between Control and Rescue crosses (*p* = 0.73). In a similar pattern, while 76% +/− 13% of Control-derived larvae and 90% +/− 11% of Rescue-derived larvae developed into adults, only 66% +/− 16% of CI-derived larvae developed into adults (Fig 5A). Adjusted pairwise comparisons demonstrated this reduction in larval survival was significant between CI-derived and both Control-derived (*p* = 0.02) and Rescue-derived (*p* = 6×10^−4^) progeny, with no significant difference between Control and Rescue-derived progeny (*p* = 0.13). The cumulative effects of these lethal phases substantially increase the strength of CI: Control egg-to-adult = 71% +/− 15% survival; CI egg-to-adult = 52% +/− 16% survival; Rescue egg-to-adult = 87% +/− 12% survival (Fig 5A). Adjusted pairwise comparisons demonstrated this reduction in overall survival was significant between CI-derived and either Control-derived (*p* = 6×10^−5^) or Rescue-derived (*p* = 1×10^−5^) progeny, with no significant difference between Control and Rescue-derived progeny (*p* = 0.16). Therefore, these studies indicate egg-to-adult assays are essential for a more accurate measure of CI strength.

**Fig 5.**
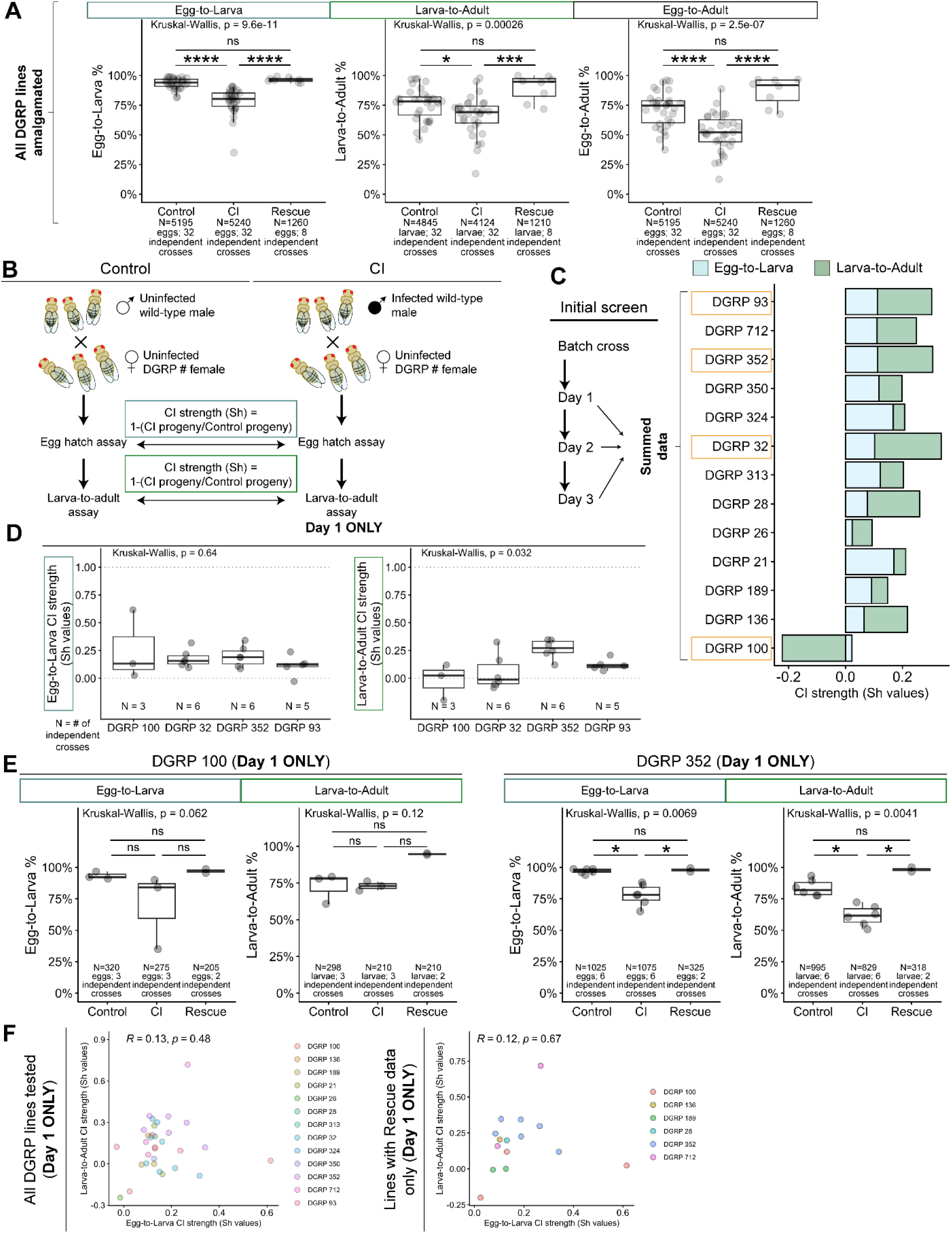
Host maternal factors influence the strength of CI-induced lethality. **A)** Graphs depicting Egg-to-Larva (left), Larva-to-Adult (middle), and Egg-to-Adult (right) survivability for Control, CI, and Rescue crosses from amalgamated DGRP experiments. **B)** Normalized pre- and post-hatching lethality was determined by calculating Sh values (1-(CI progeny/Control progeny)). **C)** For our initial screen, embryos were collected successively over three nights and Sh values for Egg-to-Larva (blue) and Larva-to-Adult (green) were determined based on summed data collected over the three days. Boxed DGRP lines indicate those chosen for further analysis. **D)** Graphs depicting Egg-to-Larva (left) and Larva-to-Adult (right) Sh values for CI crosses for each selected DGRP line, calculated from only Day 1 collections (when CI is strongest). **E)** Graphs depicting Egg-to-Larva and Larva-to-Adult survivability for crosses in which female flies were DGRP 100 (left) or DGRP 352 (right), the lines with which it was possible to perform Rescue crosses. **F)** Analysis of pre- (X-axis) and post-hatching (Y-axis) lethality phases showed little-to-no correlation in the strength of the CI-induced lethality as measured with Sh values for all DGRP lines assayed (left) and for just those for which Rescue crosses could be performed (right). Statistical significance was assessed by Dunn’s tests using the Bonferroni correction to adjust for multiple comparisons **(A,E)** **** = *p* < 0.0001; *** = *p* < 0.001; * = *p* < 0.05; ns = *p* > 0.05. Sample size data for eggs and larvae can be found in supplemental table 1 **(C,D,F)**.

### The strength of CI-induced reductions in pre- and post-hatch survivability are independent

To determine and compare pre-embryonic and post-embryonic CI strengths, we calculated the Sh value (1 – ratio of CI progeny recovered to Control progeny recovered) (7, 34) for each DGRP cross (Fig 5B). For our initial screen, we collected progeny from the same cross over three subsequent days and grouped these data together (Fig 5C). This enabled us to determine which lines to use for subsequent studies. Given their relatively high Sh values for both egg-to-larva and larva-to-adult CI strengths, we were particularly interested in crosses using lines DGRP 93, DGRP 352, and DGRP 32 (Fig 5C). Given the weak and opposing nature of the Sh values for DGRP 100, we also selected this line for further study (Fig 5C). We subsequently performed additional assays on these four lines. Because CI strength reduces with time (34, 35), we limited our following analyses to progeny collected on the first collection day (when CI should be strongest). We next determined if CI strength scores differed significantly across the DGRP lines at the egg-to-larva and the larva-to-adult stages (Fig 5D). While we observed no significant difference in the CI strengths across the different DGRP lines at egg-to-larval stages (*p* = 0.64), we observed a significant difference in the CI strengths across the different DGRP lines at the larva-to-adult stage (*p* = 0.032); however, adjusted pairwise comparisons showed no significant differences between individual pairs (DGRP 100-DGRP 32 *p* = 1; DGRP 100-DGRP 352 *p* = 0.09; DGRP 32-DGRP 352 *p* = 0.07; DGRP 100-DGRP 93 *p* = 1; DGRP 32-DGRP 93 *p* = 1; DGRP 352-DGRP 93 *p* = 0.5). Given the source of variability among these crosses solely arises from the varied genetic backgrounds of the DGRP females (all males were from the same OreR stock), these data suggests that host maternal contributions can modulate CI strength.

To determine if larva-to-adult lethality in these lines was due specifically to CI effects as opposed to lethality caused from crossing a given DGRP line to our lab stock, we measured larva-to-adult lethality in Rescue crosses. Of these 4 crosses, 2 (DGRP 100 and DGRP 352) had originally been infected with *Wolbachia* allowing us to perform Rescue crosses. For DGRP 100, we could not detect any CI induced effects at either the egg-to-larva (Control = 93% +/− 3%, CI = 70% +/− 30%, Rescue = 97% +/− 2%; adjusted pairwise comparisons: Control-CI *p* = 0.29, CI-Rescue *p* = 0.08, Control-Rescue *p* = 1) or larva-to-adult stages (Control = 72% +/− 10%, CI = 73% +/− 3%, Rescue = 95% +/− 1%; adjusted pairwise comparisons: Control-CI *p* = 1, CI-Rescue *p* = 0.13, Control-Rescue *p* = 0.35) (Fig 5E). In contrast, for DGRP 352 we observed both egg-to-larva and larva-to-adult lethality in CI-derived progeny (egg-to-larva = 78% +/− 9%; larva-to-adult = 62% +/− 8%) compared to Control-derived progeny (egg-to-larva = 97% +/− 2%; larva-to-adult = 84% +/− 6%) at a significant level (adjusted Control-CI pairwise comparisons: egg-to-larva *p* = 0.02, larva-to-adult *p* = 0.04) (Fig 5E). Importantly, Rescue-derived progeny exhibited high rates of both egg-to-larva (98% +/− 2%) and larva-to-adult (98% +/− 0.02%) survival (Fig 5E). This survivability was significantly increased compared to CI (adjusted CI-Rescue pairwise comparisons: egg-to-larva *p* = 0.04, larva-to-adult *p* = 0.01) and comparable to Control survivability (adjusted Control-Rescue pairwise comparisons: egg-to-larva *p* = 1, larva-to-adult *p* = 0.72). Thus, the observed post-embryonic lethality is due to CI and not due to hybrid inviability from crossing distinct genotypes.

To determine the relationship between pre-embryonic and post-embryonic CI-induced lethality, we compared calculated Sh values of CI strength between egg-to-larva and larva-to-adult stages (Fig 5F). Pearson coefficient analysis revealed little-to-no correlation (*R* = 0.13, *p* = 0.48) between the strength of CI-induced pre- and post-hatching lethality, even when restricting to lines for which we had performed Rescue crosses (*R* = 0.12, *p* = 0.67). These results suggest that independent factors influence the extent of pre-and post-hatching CI-induced lethality.

## DISCUSSION

The majority of published work on CI-induced phenotypes in *D. melanogaster, D. simulans*, and other species has focused on embryonic lethality prior to egg hatching. The proximal cause of this lethality is disruptions in paternal chromosome condensation and segregation during the first zygotic division. These segregation defects result in either an arrest of embryo development or haploid development followed by pre-hatching arrest (13, 36, 37, 38). However, progeny from CI-crosses also exhibit post-hatching lethality (27, 28, 39). Our recent work in *D. simulans* revealed that the CI-induced late larval lethality is distinct from and independent of the CI-induced first division defects (28). Approximately a third of CI-derived embryos that successfully hatch subsequently exhibit a post-hatching lethal phase, likely during the larval stages. Significantly, in the Rescue crosses, both pre- and post-hatching lethality are suppressed. Thus, in addition to the well documented first division segregation defects in *D. simulans* that cause pre-hatching lethality, CI also results in a distinct lethal phase later in development. Support for this interpretation comes from documentation of CI-induced post-hatching lethality in haplodiploid thrips (27).

A major goal of the work presented here was to determine whether CI produced similarly distinct lethal phases in *D. melanogaster.* We were particularly interested in examining this issue in genetically varied wild-type backgrounds as an additional lethal phase has implications for understanding the rate of spread and distribution of *Wolbachia* in wild populations. With this in mind, we took advantage of several isogenic *D. melanogaster* DGRP lines isolated from nature (31). In accord with previous reports, we observed a significant but modest reduction in egg hatch in embryos derived from CI crosses (that is, much less pronounced than observed in *D. simulans*). We also observed a significant reduction in larva-to-adult survival in progeny derived from the *D. melanogaster* CI crosses. This CI-induced post-hatching lethality was eliminated in the Rescue cross. Given the first division segregation errors result in pre-hatching lethality and generally weaker first division defects observed in *D. melanogaster* overall, this CI-induced post-hatching lethality is unlikely due to failed development resulting from first division defects. Consistent with this, the strength of post-hatching lethality is not correlated with the strength of the first division defect as determined by measuring pre-hatching lethality. This suggests the CI-induced pre- and post-hatching lethality are at least in part regulated by distinct factors (Fig 6A). Collectively, these results reveal that in both *Drosophila* species, CI has a more pronounced lethality than previously documented in studies focused only on egg hatch.

**Fig 6.**
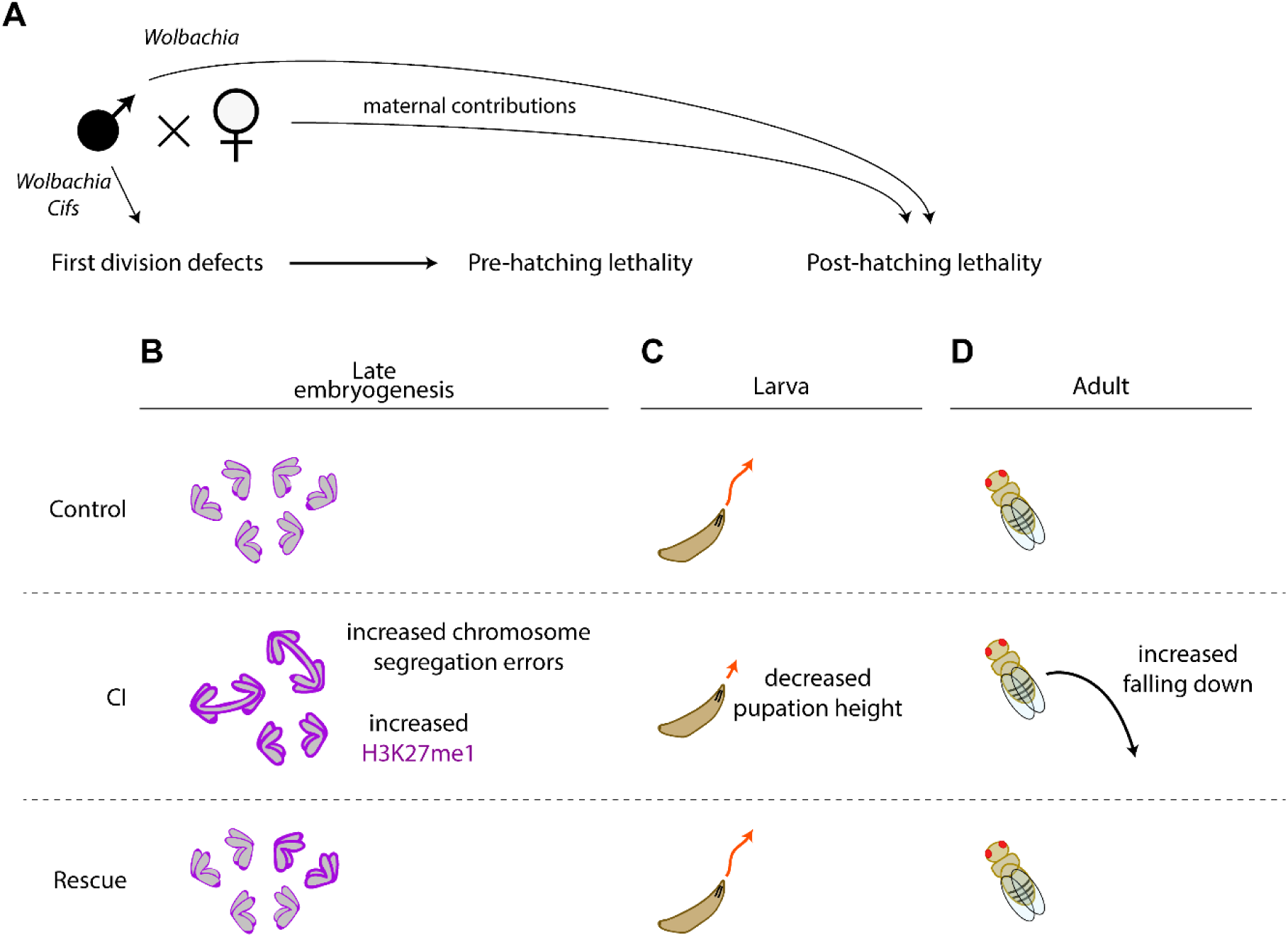
Both *Wolbachia* and host maternal factors influence the strength of CI-induced lethality. **A)** CI-induced pre-hatching lethality is due to the action of *Wolbachia*-produced *Cifs* that cause first division chromosome segregation defects. In addition, there is a second post-hatching lethal phase independent from the first division defects. Maternal contributions influence the strength of the post-hatching lethal phase. Whether maternal contributions contribute to the other CI-induced phenotypes has yet to be determined **B)** Gastrulating CI-derived embryos experience higher rates of chromosome segregation errors and levels of the H3K27me1 chromatin mark. **C)** CI-derived larvae experience locomotor defects and reduced pupation height which are restored in Rescue crosses. **D)** CI-derived adults experience increased falling which is rescued in the Rescue cross.

Additionally, although CI has been extensively investigated in *D. melanogaster*, surprisingly few studies have been performed examining the embryonic cytological effects in naturally infected strains. Studies of *D. simulans* CI-derived embryos reveal extensive division errors during the late cortical divisions, cellularization, and in the mitotic patches of gastrulating embryos (28). This suggests that the *Wolbachia*-induced perturbations of the paternal chromosome complement are inherited and preserved through multiple rounds of mitotic division. As discussed below, potential insight into this phenomenon comes from our finding that CI induces dramatic increases in embryonic H3K27 mono-methylation levels (Fig 6B). It is unclear why segregation errors manifest during gastrulation and not earlier. Significant developmental events at this time include the establishment of a G2 cell cycle phase, facultative heterochromatin formation, and the transition from maternal to zygotic transcriptional regulation (40, 41, 42). These events may stress already sensitized chromatin. Thus CI-induced disruptions to the chromatin may be responsible for the segregation defects.

To determine if *D. melanogaster* manifests additional CI-induced developmentally delayed phenotypes, we examined the larval stages. While there were no obvious abnormalities in larval development or morphology, we discovered that the 3rd instar larvae pupate at a much lower height than Control larvae. This indicates that the climbing ability of the third instar larvae is compromised (Fig 6C). Support for the idea that this defect may have a neurological basis, comes from previous studies that have shown a broad spectrum of neuronal mutants, including *scribbled*, a gene involved in nervous system development, result in reduced pupation height (43). While these climbing defects may not affect adult survival under ideal lab conditions, they may be consequential in natural settings. Larvae that fail to climb out of the rotting fruit are more susceptible to mold, predators and hypoxia (44). Indeed, we observed many of the CI-derived larvae were unable to climb out of the food plate even in a lab environment. In addition, robust climbers are better able to leave crowded conditions and disperse to preferred pupation sites (45). Finally, these locomotor defects continue onto adulthood with CI-derived adults more likely to fall off the vial wall while climbing (Fig 6D).

These studies provide baseline wild-type data for future genetic studies identifying host factors that influence CI strength. While environmental factors such as temperature, age, developmental timing, and diet (35, 46, 47, 48) are known to potentiate CI, key studies have only just begun to identify host genetic factors that influence CI strength (49, 50, 51, 52). By taking advantage of the genetic variation in DGRP isolines, our results indicate that maternally derived factors play an important role in determining the strength of CI (Fig 6A). Uninfected females from thirteen DGRP lines were crossed to the same stock of *w*Mel-infected OreR males and egg hatch rates and larva-to-adult survival rates of the progeny were determined. These studies revealed extensive variation between isolines with respect to larva-to-adult survival, with no significant correlation between the pre-embryonic and post-embryonic lethal phases. This suggests distinct factors, including host factors, influence the strength of these two lethal phases. Whether the variation is due to maternal effect factors or maternally derived zygotic factors in unknown. In addition, as we did not control for any differences in gut microbiota between the lines, it is possible that gut microbiota may be a contributing factor in these experiments. Because these studies suggest that genetic variation in females results in tremendous variation in CI strength, one would suspect in predominantly uninfected populations there would be strong selection for those females bearing modifiers that reduce CI-induced lethality.

The identity of the host factors that influence CI strength remain to be determined. Potential insight comes from our finding that CI induces an increase in embryonic H3K27me1 levels (Fig 6B), consistent with results in *Nasonia* (30). However, we note that the methylation levels were extremely variable, with a notable overlap in H3K27me1/DAPI ratios between Control-, CI-, and Rescue-derived gastrulating embryos. Therefore, we urge caution in interpreting a causative link between increased methylation and subsequent CI-induced defects. Nevertheless, increased H3K27 methylation levels in *Drosophila* CI-derived embryos suggest genes regulating the state of the host chromatin may be involved. Interestingly, a number of diverse endosymbionts produce effector proteins termed “nucleomodulins” that are capable of remodeling host chromatin (53). It may be that host modifiers counter the action of potential *Wolbachia*-produced nucleomodulins. We suspect some of these modifiers are either the target of or interact with the targets of *Cifs*, the *Wolbachia* cytoplasmic incompatibility factors (10).

## MATERIALS AND METHODS

### Drosophila stocks

All fly stocks were grown on standard brown food (a mix of roughly 80% water, 8% molasses, 0.7% agar, 6% cornmeal, 2.5% yeast, 0.13% Tegosept 10%, 1% Ethanol, and 0.5% propionic acid) (54) at 25°C with a 12 hr light/dark cycle in a Bally BSC walk-in incubator (model number: 3478). The flies used in this study include *w*Mel-infected and tetracycline-cured Oregon-R (OreR) wild-type *Drosophila melanogaster* maintained in the lab for over 10 years. (Figs 2-5) (55, 56). We used several *Drosophila* Genetic Reference Panel (DGRP) lines (31) obtained from the Bloomington *Drosophila* Stock Center (Fig 5). These stocks include the following DGRP lines:*21*(RRID:BDSC_28122), *26*(RRID:BDSC_28123), 28(RRID:BDSC_28124), *32*(RRID:BDSC_55015), *93*(RRID:BDSC_28137),*100*(RRID:BDSC_55017),136(RRID:BDSC_2814), *189*(RRID:BDSC_28152),*313*(RRID:BDSC_25180,*324*(RRID:BDSC_2518, 350(RRID:BDSC_28176).*352*(RRID:BDSC_83728),*712*(RRID:BDSC_252).

Each DGRP line was initially assessed for *Wolbachia* infection by PCR and gel electrophoresis using primers amplifying the 16s rRNA *Wolbachia* gene (57) (Forward: TCATGTACTCGAGTTGCAGAGT, Reverse: ATACGGAGAGGGCTAGCGTTA).

Infected stocks (DGRP 28, 93, 100, 136, 189, 352, and 712) were split with one subline maintained as an infected stock, while the other was treated with tetracycline for 3 generations to generate a corresponding uninfected stock (58). All *Drosophila* stocks were routinely assessed by PCR and gel electrophoresis for *Wolbachia* infection status. Cured DGRP stocks were used immediately after a negative PCR result was obtained.

### Egg hatch (egg-to-larva) assays

For Control, CI and Rescue crosses, uninfected and infected virgins were collected over a period of 2 days. Virgin flies were identified via the meconium spot. Virgin flies were anesthetized for at most 15 to 20 minutes using CO₂. The females and males were sorted by sex via their genitals and segregated into standard brown food vials for 1 to 2 days. Uninfected and infected virgin females were mated to either infected or uninfected unmated 1-2 day old males according to the cross. Each cross consisted of approximately 25-50 females and an equal number of males. For crossing, flies were anesthetized with CO2, then transferred to a collection bottle (Fisherbrand round Drosophila bottles with a standard brown food plate (3.5 cm Corning plastic petri dish) attached to the mouth of the bottle). For experiments involving DGRP lines, DGRP females were collected in the same manner as described above and crossed to wild-type OreR *w*Mel infected or uninfected *D. melanogaster* males that were collected as described above.

For Fig 5C we collected eggs from 5pm to 9am over 3 sequential nights after an initial 24-hour mating period. For all other experiments eggs were collected only during the first 5pm to 9am period following the initial 24-hour mating period. Collected eggs were transferred to a new brown food plate and incubated at 25°C for 48 hours after which the number of unhatched eggs were counted.

DGRP egg hatch assays were not performed simultaneously but over the same period of time. Then assays with *OreR* lab stocks were performed after. Control, CI, and Rescue crosses were all performed simultaneously side by side for all experiments.

### Larva-to-adult assays

Larva-to-adult assays were performed with the larvae that emerged from the egg hatch assays. The food plates that were used in the egg hatch assays were placed onto empty collection bottle and were kept for 10 days at 25°C. The number of emerging adults were counted over a period of 3 days after the first adult emergence. The bottle was maintained for an additional 24 hours to ensure that no additional adult flies emerged.

DGRP larva-to-adult assays were not performed simultaneously but over the same period of time. Then assays with *OreR* lab stocks were performed after. Control, CI, and Rescue crosses were all performed simultaneously for all experiments.

### Pupation height assay

1-2 day old female and male virgins were crossed (Control, CI and Rescue crosses) as described above. We collected eggs from both 1) 5pm to 9am over 3 sequential nights after an initial 24-hour mating period and 2) only during the first 5pm to 9am period following the 24-hour mating period (18 collections were performed from 8 independent crosses). Collected eggs were transferred to a new brown food plate and an egg hatch assay was performed as described above. 7 days after the egg hatch was determined, pupae both on the bottle and in the food plate were counted and the number of pupae above/below a 2 cm line (the physical rim at the neck of the bottle) were recorded.

### Negative-geotaxis assay (adult climbing)

Negative-geotaxis assays were performed to assess potential locomotor defects. We used the same crossing and rearing protocol described from egg hatch and larva-to-adult assays; however, these experiments were conducted independently using separate cohorts of lab stocks. Adult male and female flies were collected from Control, CI, and Rescue crosses by anesthetizing with CO₂ for no longer than 15 to 20 minutes and sorted into groups of 25 individuals per each cross. Flies were maintained in standard brown food vials for 24 hours at 25°C prior to testing.

For the assay, flies were transferred without anesthetizing to a clean, empty chamber, consisting of 2 vials taped together, mouth to mouth, marked with a line 15 cm from the bottom. Immediately before recording, flies were gently tapped to the bottom of the vial to initiate the climbing response. A DinoLite digital camera was positioned so that the 15 cm line was centered in the field of view, and the flies were recorded for 1 minute (15 fps). The number of flies crossing the 15 cm line in either an upward or downward motion was recorded.

Each vial of flies was assayed three times per trial, and 7 independent biological replicates were performed using separate cohorts of adults derived from the independent crosses. At 10-second intervals, 3 measurements were recorded: 1) the number of flies that crossed above the 15 cm line, 2) the number of flies that subsequently fell below the 15 cm line, and 3) the number of flies that crawled below the 15 cm line. These three measurements enabled us to avoid double counting flies that crossed the line more than once. Downward movement was further categorized to distinguish between flies that fell and those that actively descended by crawling. Falling events were identified by rapid, uncontrolled downward movement through the vial, allowing them to be distinguished from active flight or voluntary downward crawling.

In total 175 flies were tested for Control-, CI-, and Rescue-derived adults. To estimate the rate of falling the number of flies that had fallen was divided by the number of flies that had made it above the 15 cm line.

### Embryo fixation

For experiments assaying embryo chromosome segregation abnormalities (Fig 2), OreR male and female flies were crossed following the same procedure as described in “Egg hatch assays.” Females were allowed to lay for 2 hours, and those eggs were collected and transferred to a fresh food plate to age for an additional 3 hours at 25°C. These 3-5 hour old embryos were then dechorionated in 25% bleach, washed thoroughly in water, and transferred to 1 mL of ice-cold heptane for 2.5 minutes. 1 mL of ice-cold methanol was added and embryos were shaken vigorously for approximately 1 minute (54). Heptane was removed and embryos were stored in fresh methanol at 4°C overnight. Embryos were mounted the following day, directly in DAPI with Vectashield (Vector H-1200-10).

For experiments analyzing methylation (Fig 3), 3-5 hour embryos were initially fixed via boiling fixation (54): embryos were dechorionated with 25% bleach, rinsed thoroughly with water, and placed in a 50 mL conical tube containing 5 mL of embryo wash solution (0.7% NaCl and 0.05% Triton X-100) heated to ∼90-100°C. 40 mL of ice-cold embryo wash was added immediately. Embryos were then transferred to a 5 mL glass vial for devitellinization with 1 mL of heptane followed by 1 mL of methanol. The vial was shaken vigorously for 15 seconds. Heptane and methanol layers were removed, and 4 mL of fresh methanol was added to the vial. Embryos were stored at 4°C for 24 hours. Embryos were rehydrated in PBT (phosphate-buffered saline (PBS) + 0.05% Triton X-100), blocked for 1 hour in PBT containing 1% of bovine serum albumin (BSA) (Roche 10735086001). Embryos were incubated overnight at 4°C with a rabbit anti-H3K27me1 polyclonal antibody (1:1000 Active Motif 39378). After removing the primary antibody, embryos were washed 3 times for 20 minutes in PBT, embryos were incubated for an hour at room temperature with anti-rabbit-Alexa488 secondary (1:500 Thermo Fisher A-11008). After removing the secondary antibody, embryos were washed 3 times in PBT for 20-minute washes, rinsed 2× in PBS, and mounted with DAPI in Vectashield (Vector Laboratories H-1200-10).

3-5 hour embryos in the gastrula stage (as determined by classic features of gastrulae such as cephalic furrow formation and distinct mitotic domains) were specifically imaged.

### Confocal microscopy

Fixed embryo imaging was performed on a Leica Stellaris confocal microscope. DAPI was excited with a 405 nm laser and collected from 410 to 480 nm. Alexa488 was excited with a 488 nm laser and collected from 518 to 584 nm. Embryos were imaged with either 10×/0.3 dry objective or 63×/1.4 oil objective. All imaging was performed at room temperature. Images were acquired with the Stellaris Leica LAS X software.

No deconvolution was performed. Brightness and contrast were adjusted for clarity in FIJI. All qualifications were performed on raw, unadjusted images.

### Quantification and statistical analysis

Given the hierarchical nature of some our data that involved multiple offspring analyzed from batch crosses (Figs 2D, 3E) or multiple collections from the same cross (4C-D) or multiple measurements of the same flies (4G-H), we generated linear and generalized linear mixed models using the “lmer” function of the lme4 package or the “glmmTMB” function (3E, family=Gamma(link=”log”); 4G, family=beta) of the glmmTMB package in R (59, 60). We set the genotype as the fixed effect and the batch, collection, or measurement as the random effect. Models were assessed with the DHARMa and Performance packages (61, 62) and adjusted to improve the fit of residuals. Degrees of freedom for linear mixed models were calculated using the Kenward-Roger method. For normalized H3K27me1/DAPI ratios (Fig 3E), while assessment of the generated model using the DHARMa package showed significant within-group deviations from uniformity for the model, there was no significant over- or under-dispersion (non-parametric dispersion test, *p* = 0.63), and residuals were uniformly distributed (Kolmogorov-Smirnov test, *p* = 0.11). Therefore, we interpret the within-group variation to represent minor sample variations instead of a major fault of the model. Pairwise comparison tests were performed to compare differences between estimated marginal means for Control, CI, and Rescue crosses using the emmeans package (63), applying the Tukey method for *p* value adjustment for multiple comparisons. 95% confidence intervals (95%CI) and *p* values of comparisons are reported.

All other egg-to-larva hatch rate (Fig 2A, Fig 4I, Fig 5A, E), larva-to-adult (Fig 5A, E), and egg-to-adult (Fig 5A), comparisons were made with Kruskal-Wallis tests and subsequent Dunn’s tests, applying the Bonferroni method for *p* value adjustments for multiple comparisons. Variation of CI strength across different maternal DGRP lines (Fig 5D) was determined with Kruskal-Wallis tests.

Possible correlation between pre- and post-hatching lethality (Fig 5E) was assessed by calculating the Pearson correlation coefficient.

Reported +/−% values throughout the results section represent the standard deviations among replicates.

All statistics were performed in R (version 4.5.2).

### Figure preparation

Graphs were created in R using the ggplot2 package (64). To improve the clarity of certain panels, images were adjusted for brightness and contrast in FIJI. Figures were assembled in Adobe Illustrator (Adobe, San Jose, CA, USA). Data tables were created in Microsoft Word (Microsoft, Redmond, WA).

## Supporting information

raw data

## ACKNOWLEDGEMENTS

We thank Benjamin Abrams (UCSC Life Sciences Microscopy Center, RRID: SCR_021135) for his technical support and microscopy assistance. We thank Michael Grant for support and assistance. Funding for these studies was provided by National Institutes of Health grant NIGMS-5R35GM139595 awarded to W.S.

## DECLARATION OF INTERESTS

The authors declare no competing interests.

## AUTHOR CONTRIBUTIONS

Conceptualization: C.P., J.P., W.S.; Methodology: C.P., J.P., B.W., W.S.; Investigation: C.P., J.P., B.W., W.S.; Validation: C.P., J.P., B.W., W.S.; Formal Analysis: C.P., J.P., B.W., W.S.; Resources: W.S.; Data Curation: C.P., J.P., B.W., W.S.; Writing-Original Draft: C.P., J.P., B.W., W.S.; Writing-Review & Editing: C.P., J.P., B.W., W.S.; Visualization: C.P., J.P., B.W., W.S.; Supervision: B.W., W.S.; Project Administration: W.S.; Funding Acquisition: W.S.

---

**S1 Table.** Sample sizes of eggs and larvae data corresponding to figure 5 C,D, and F.

|  | Control |  | CI |  |
| --- | --- | --- | --- | --- |
|  | Eggs | Larvae | Eggs | Larvae |
| DGRP-93 | 545 | 494 | 600 | 483 |
| DGRP-712 | 675 | 603 | 665 | 528 |
| DGRP-352 | 575 | 522 | 600 | 483 |
| DGRP-350 | 620 | 554 | 590 | 465 |
| DGRP-324 | 560 | 493 | 620 | 454 |
| DGRP-32 | 450 | 402 | 450 | 361 |
| DGRP-313 | 600 | 534 | 585 | 457 |
| DGRP-28 | 550 | 402 | 550 | 371 |
| DGRP-26 | 555 | 513 | 555 | 501 |
| DGRP-21 | 600 | 552 | 550 | 420 |
| DGRP-189 | 545 | 496 | 600 | 496 |
| DGRP-136 | 600 | 582 | 600 | 544 |
| DGRP-100 | 505 | 464 | 440 | 395 |

**Figure 5D**
|  | Control |  | CI |  |
| --- | --- | --- | --- | --- |
|  | Eggs | Larvae | Eggs | Larvae |
| DGRP-93 | 765 | 691 | 905 | 724 |
| DGRP-352 | 1025 | 995 | 1075 | 829 |
| DGRP-32 | 860 | 807 | 820 | 596 |
| DGRP-100 | 320 | 298 | 275 | 210 |

**Figure 5F**
|  | Control |  | CI |  |
| --- | --- | --- | --- | --- |
|  | Eggs | Larvae | Eggs | Larvae |
| DGRP-93 | 765 | 691 | 905 | 724 |
| DGRP-712 | 425 | 404 | 400 | 311 |
| DGRP-352 | 1025 | 995 | 1075 | 829 |
| DGRP-350 | 205 | 182 | 200 | 155 |
| DGRP-324 | 210 | 182 | 200 | 149 |
| DGRP-32 | 860 | 807 | 820 | 596 |
| DGRP-313 | 200 | 178 | 185 | 146 |
| DGRP-28 | 150 | 122 | 150 | 106 |
| DGRP-26 | 155 | 139 | 155 | 141 |
| DGRP-21 | 375 | 362 | 325 | 269 |
| DGRP-189 | 305 | 290 | 350 | 297 |
| DGRP-136 | 200 | 195 | 200 | 175 |
| DGRP-100 | 320 | 298 | 275 | 210 |

## Notes

### Competing Interest Statement

The authors have declared no competing interest.

